# Isolation of full-length IgG antibodies from combinatorial libraries expressed in the cytoplasm of *Escherichia coli*

**DOI:** 10.1101/2020.05.09.085944

**Authors:** Michael-Paul Robinson, Emily C. Cox, Mingji Li, Thapakorn Jaroentomeechai, Xiaolu Zheng, Matthew Chang, Mehmet Berkmen, Matthew P. DeLisa

**Author notes:** Address correspondence to: Matthew P. DeLisa, Robert Frederick Smith School of Chemical and Biomolecular Engineering, Cornell University, Ithaca, NY 14853.

## Abstract

We describe a facile and robust genetic selection for isolating full-length IgG antibodies from combinatorial libraries expressed in the cytoplasm of the genetically engineered *Escherichia coli* strain, SHuffle. The method is based on the transport of a bifunctional substrate comprised of an antigen fused to chloramphenicol acetyltransferase, which allows positive selection of bacterial cells co-expressing cytoplasmic IgGs called ‘cyclonals’ that specifically capture the chimeric antigen and sequester the antibiotic resistance marker in the cytoplasm. The selective power of this approach was demonstrated by facile isolation of novel complementarity-determining regions for a cyclonal that specifically recognized the basic-region leucine zipper domain of the yeast transcriptional activator protein Gcn4.

## Introduction

Monoclonal antibodies (mAbs) represent one of the fastest growing segments of the biotechnology industry, enabling dramatic advances in biomedical research and modern medicine. Consequently, new technologies capable of straightforward identification and molecular engineering of full-length immunoglobulin G (IgG) antibodies are in great demand. Conventional procedures for isolating mAbs include various hybridoma technologies that involve immunization followed by cell fusion ^1-3^ and protein engineering platforms such as phage display ^4^, ribosome and mRNA display ^5^, and microbial cell display technologies ^6-9^ that permit high-throughput screening of large recombinant antibody libraries. Hybridoma-based methods for isolating antibodies are labor- and time-intensive, incompatible with multiplexing and parallelization, and do not permit customization of mAb properties such as antigen-binding affinity, stability, or expression level. These shortcomings can be circumvented by the use of display technologies; however, these methods are typically built around libraries of smaller, more conveniently expressed derivatives of mAbs such as single-chain variable antibody fragments (scFv) or antigen-binding fragments (Fabs) ^8^. Compared to their IgG counterparts, these smaller formats exhibit weaker monovalent binding and poor serum persistence in animals, the latter of which stems from their relatively low molecular weight and lack of an Fc domain. Consequently, antibody fragments isolated using display technologies require molecular conversion to IgG format prior to therapeutic development.

More recently, cell surface display of full-length IgGs has been demonstrated in bacteria ^10-12^, yeast ^13-15^, and mammalian cells ^16^, effectively circumventing the reformatting issue. Nonetheless, screening methods such as these require each library member to be individually evaluated, which necessitates specialized equipment (*e*.*g*., flow cytometer) to access meaningful amounts of sequence space and a high-quality screening antigen, typically a purified recombinant protein that must be separately prepared. It should be noted that even with state-of-the-art instrumentation, the screening of combinatorial libraries with diversity >10^8^ is technically challenging ^12,17^. Another drawback of cell surface display is the inherent bias and complexity that can be introduced by the need for energetically unfavorable trafficking of IgG molecules across one or more biological membranes, which are known to selectively eliminate clones that are unfit for translocation but might otherwise be viable. Moreover, IgG display in yeast cells involves a secretion-capture process that is prone to “crosstalk” among library members while in mammalian cells the process suffers from limited library sizes due to low transfection efficiency and the appearance of multiple copies of antibodies with different specificities on a single cell surface, making it difficult to identify and isolate antibodies with desired properties from naïve libraries.

To address these shortcomings, here we describe a genetic selection strategy for isolating full-length IgGs from combinatorial libraries expressed in the cytoplasm of *Escherichia coli*. Specifically, the method leverages bifunctional substrate proteins comprised of an antigen fused to chloramphenicol acetyltransferase (CAT) whose translocation through the twin-arginine translocation (Tat) pathway ^18^ is toggled by the presence or absence of a cytoplasmically co-expressed IgG antibody known as a ‘cyclonal’ ^19^. In this manner, capture of the chimeric antigen by a co-expressed IgG effectively sequesters the CAT antibiotic resistance marker in the cytoplasm, leading to detoxification of chloramphenicol and permitting positive selection for antigen binding. By using the genetically engineered *E. coli* strain SHuffle, which promotes efficient cytoplasmic disulfide bond formation ^20^, it is possible to achieve high-level functional expression of cyclonals within the cytoplasmic compartment while at the same time bypassing the need for membrane translocation of IgG molecules. Moreover, compared with screening methods that necessitate analysis of each individual IgG variant, our selection directly eliminates unwanted IgG variants through the application of tunable selective pressure on the mutant library. This feature of selection makes it intrinsically high throughput, enabling assessment in theory of very large libraries (>10^11^). We demonstrated the utility of this approach by isolating cyclonals with novel complementarity-determining regions (CDRs) that promote specific binding to the basic-region leucine zipper domain of the yeast transcriptional activator Gcn4. Importantly, discovery of these CDR variants was made possible by simply demanding bacterial growth on defined concentrations of antibiotic, obviating the need for purification or immobilization of the target antigen. Hence, our selection represents a straightforward tool for enrichment of productive binders in the IgG format and offers a compelling alternative to conventional methods that are more expensive, time-consuming, and labor-intensive.

## Results

### Design of a positive selection for antigen-binding activity in the cytoplasm

The principle of our selection scheme for detecting antigen-binding activity in the cytoplasm of *E. coli* is illustrated in **Fig. 1a**. It involves the creation of a chimeric antigen biosensor in which the peptide or protein target of an antibody is genetically fused to the C-terminus of the CAT antibiotic resistance protein, which itself is modified at its N-terminus with the signal peptide derived from *E. coli* trimethylamine *N-*oxide reductase (spTorA) that is well known to deliver completely folded guest proteins into the periplasmic compartment via the Tat export pathway ^21,22^. The rationale for the design came from previous studies demonstrating the exportability of CAT when fused to Tat signal peptides ^23^ as well as the use of CAT as a reliable genetic reporter of Tat export ^24^.

**Figure 1.**
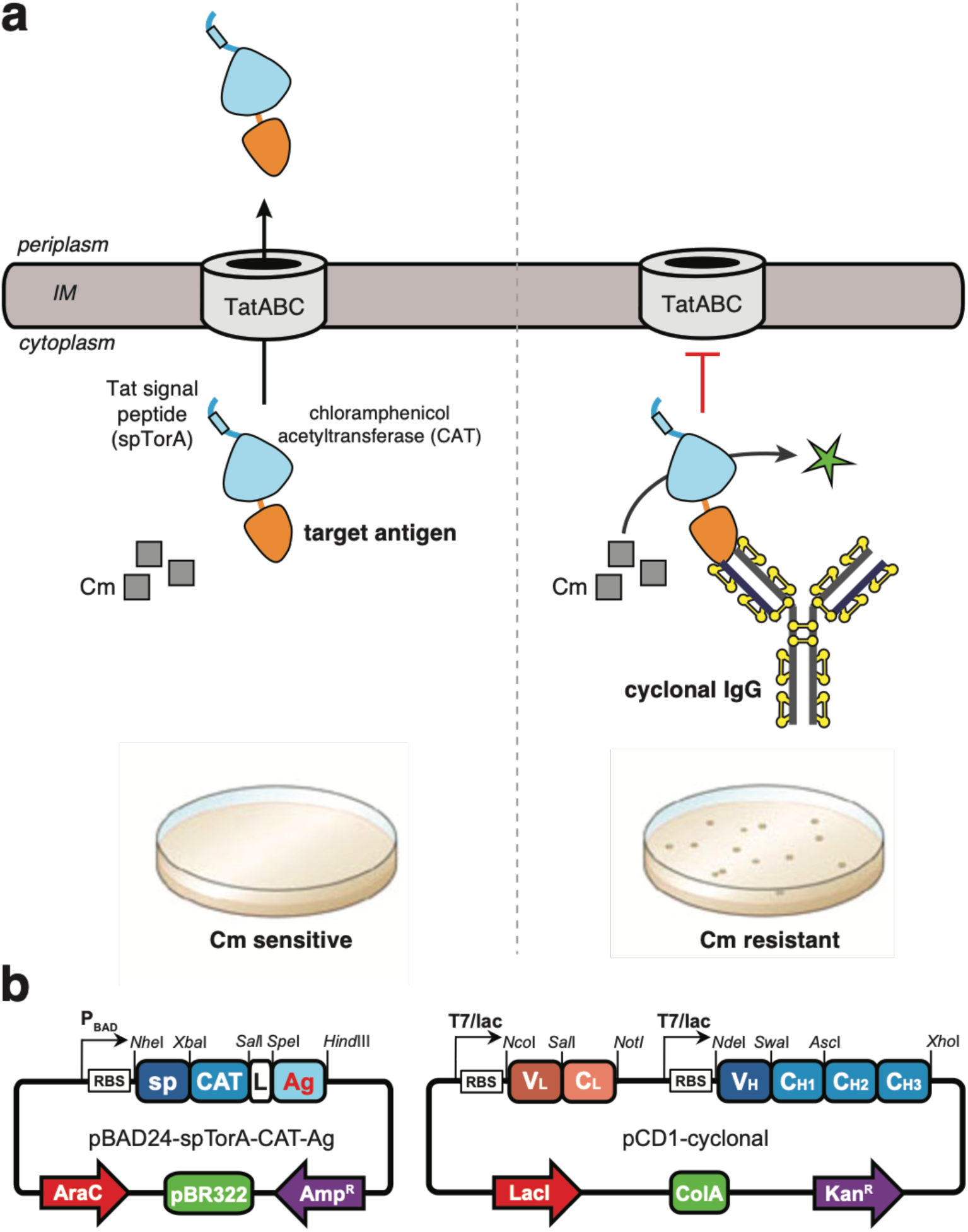
Schematic of Tat-dependent positive selection for antigen-binding activity of full-length IgGs. In the absence of a cognate binding protein, a chimeric antigen comprised of a peptide or protein antigen of interest (orange) fused to the C-terminus of spTorA-CAT (blue) is exported out of the cytoplasm by the TatABC translocase. By localizing the chimeric antigen into the periplasm, the CAT domain is no longer able to inactivate the antibiotic chloramphenicol (Cm) by acetylation using acetyl-coenzyme A (CoA) and the host cells are rendered sensitive to antibiotic. When a cyclonal (yellow/black) is functionally expressed in the cytoplasm of SHuffle T7 Express cells, it binds specifically to the chimeric antigen, thereby sequestering the CAT domain in the cytoplasm where it can efficiently detoxify Cm (green star) and conferring an antibiotic-resistant phenotype. Individual clones from the selection plate are selected, genetically identified, and functionally characterized. Yellow balls/sticks represent the 16 intra- and intermolecular disulfide bonds in IgG that are required for folding and activity. (b) Schematic of pBAD24-based vector for expression of chimeric antigen constructs (left) and pCD1-based vector for expression of cyclonal IgGs (right). Abbreviations: RBS, ribosome-binding site; sp, TorA signal peptide; L, flexible GTSAAAG linker; Ag, antigen; V_H_, variable heavy; V_L_, variable light; CH, constant heavy; CL, constant light.

Following expression and folding in the cytoplasm, and in the absence of a cognate binding protein, the activity of the biosensor is decreased because the CAT domain of the chimeric antigen is translocated into the periplasm where it is no longer able to inactivate the antibiotic chloramphenicol by acetylation using acetyl-coenzyme A (CoA). However, if a cyclonal that binds to the peptide or protein antigen is co-expressed, it will specifically capture the chimeric antigen and sequester CAT in the cytoplasm, leading to an increase in biosensor activity due to CAT-mediated detoxification of chloramphenicol. Because the Tat system is capable of exporting multimeric protein complexes that have assembled in the cytoplasm prior to export ^25-27^, the rationale for this design is that the expected size and three-dimensional bulkiness of a cyclonal-chimeric antigen complex would exceed the capacity of the Tat system and thus be blocked for export. Indeed, the upper limit for natural *E. coli* Tat substrates is represented by the PaoA heterotrimer (MW = ∼135-kDa, radius of gyration, *R*_g_ = 35 Å, and maximal dimension, *D*_max_ = 120 Å ^28^), whereas for a human IgG1 antibody alone the size is significantly larger (150 kDa, *R*_g_ = 53 Å, *D*_max_ = 160 Å ^29^). An advantage of this approach is its simplicity as antimicrobial resistance can easily be determined in spot titer experiments, which enable the effects of mutations on protein properties (*e*.*g*., expression level, folding and assembly, binding affinity) to be phenotypically compared and quantified. Moreover, if the selectable marker is efficient enough, it should be possible to custom tailor cyclonals by selecting for variants with improved properties.

### Chimeric antigens targeted to Tat pathway inhibit cell viability

To validate our selection scheme, we first determined whether Tat export of these chimeric antigens yielded the expected chloramphenicol-sensitive phenotype. Specifically, we constructed a vector encoding a tripartite spTorA-CAT-Ag fusion where Ag corresponds to one of three different peptide antigens: (i) a 6-residue epitope from the hemagglutinin protein of influenza virus (HAG) ^30^; (ii) a 10-residue epitope from the human c-Myc proto-oncogene product ^31^; and (iii) the 47-residue basic-region leucine zipper domain of yeast Gcn4 carrying two helix-breaking proline mutations that disrupt the helical structure of the zipper and prevent its coiled-coil-mediated homodimerization (Gcn4-PP) (**Fig. 1b**) ^27,32^. Each of these epitopes was genetically fused to the C-terminus of CAT via a seven-residue flexible linker (Gly-Thr-Ser-Ala-Ala-Ala-Gly). Importantly, all three chimeric fusions were unable to confer resistance to cells that were spot plated on agar supplemented with chloramphenicol (**Fig. 2**), as we had predicted. Spot plating of the same cells on agar that lacked chloramphenicol resulted in strong growth, indicating that the inability of these constructs to confer resistance on chloramphenicol was not due to a general growth defect. To determine whether chloramphenicol sensitivity was dependent on functional Tat export of the chimeric antigens, we generated mutant versions of each construct in which the two essential arginine residues of the twin-arginine motif in the spTorA signal peptide were mutated to lysines, a substitution that is well known to completely abolish export out of the cytoplasm ^21,33^. Indeed, all spTorA(KK)-CAT-Ag chimeras were blocked for export as evidenced by the strong resistance to chloramphenicol that each of these constructs conferred to bacterial cells (**Fig. 2**). Importantly, these results indicate that the subcellular location of chimeric antigens can be discriminated by selective plating on chloramphenicol, and that cell survival depended on disrupting the Tat-dependent export of the CAT-containing chimera.

**Figure 2.**
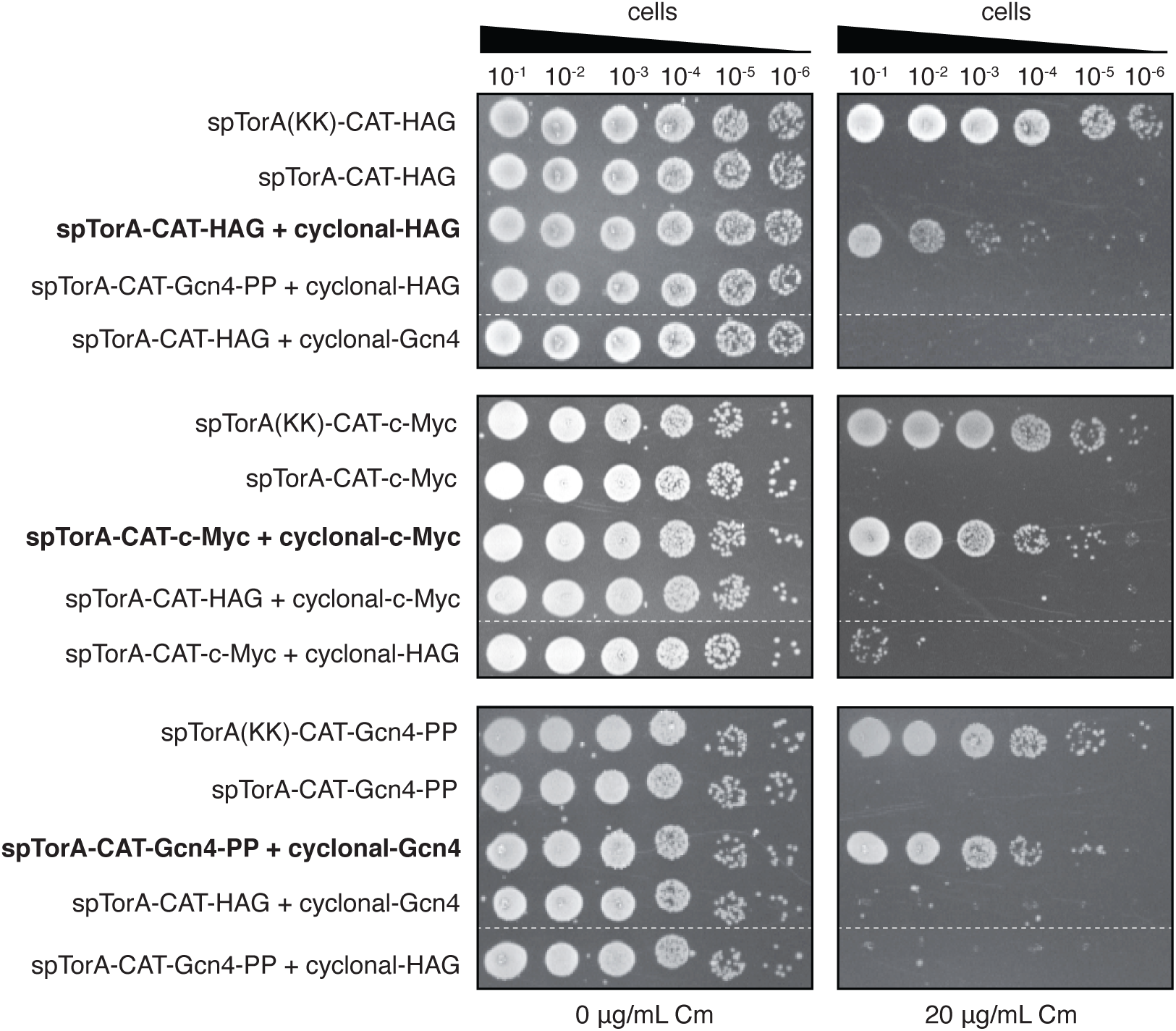
Genetic selection for cyclonal antigen-binding activity. Selective spot plating of SHuffle T7 Express cells carrying a plasmid encoding one of the chimeric antigens (spTorA-CAT-Ag or an export-defective variant spTorA(KK)-CAT-Ag) alone or with a second plasmid encoding a full-length cyclonal IgG specific for HAG, Gcn4-PP, or c-Myc as indicated at left. A total of 5 μl of 10-fold serial diluted cells was plated on LB-agar supplemented with 0 or 20 μg/ml chloramphenicol (Cm) as well as 0.4 % arabinose and 1 mM isopropyl β-D-thiogalactopyranoside (IPTG) to induce chimeric antigen and cyclonal expression, respectively. Cross-pairing the anti-HAG cyclonal with non-cognate c-Myc or Gcn4-PP and the anti-Gcn4 cyclonal with non-cognate HAG served as negative controls. Spot plating results are representative of at least three biological replicates. Dashed white lines indicate spot plating data merged from discontinuous region of plate.

### Cyclonal expression rescues cell growth in an antigen-specific manner

To determine whether antimicrobial resistance could be positively linked to the antigen-binding activity of full-length IgGs, cyclonals specific for the HAG, c-Myc and Gcn4-PP epitopes were co-expressed with their cognate chimeric antigens. Specifically, genetically engineered SHuffle T7 Express cells, which facilitate efficient cytoplasmic disulfide bond formation ^20^, were co-transformed with a plasmid encoding the cyclonal synthetic heavy and light chains, each lacking canonical export signals (**Fig. 1b**), along with a plasmid encoding the cognate chimeric antigen. When these cells were spot plated on agar supplemented with chloramphenicol, a clear increase in resistance was observed that was on par with the resistance conferred by the spTorA(KK)-CAT-Ag constructs expressed alone (**Fig. 2**). To determine whether this resistance phenotype was dependent on specific recognition of the epitope, each of the chimeric antigens was co-expressed with a non-cognate cyclonal (*e*.*g*., anti-HAG cyclonal cross-paired with spTorA-CAT-c-Myc). In all cases, there was little to no observable resistance for any of the control combinations tested, indicating that the observed antibiotic resistance was governed by antigen specificity. As above, cells grown in the absence of chloramphenicol grew robustly, indicating that cytoplasmic co-expression of these constructs had no apparent effect on cell viability. Taken together, these results unequivocally demonstrate that cyclonal IgGs sequester only their cognate chimeric antigens in the cytoplasm and significantly increase resistance by protecting cells from chloramphenicol toxicity.

We also tested whether the genetic selection presented here could discriminate the binding activity of different cyclonal variants. For this experiment, we focused on the anti-Gcn4 cyclonal because previous studies identified a number of mutations within the 5-residue heavy-chain CDR3 (CDR-H3) of single-chain Fv intrabodies whose binding activity was quantified *in vivo* and *in vitro* ^27,32^. Starting with the parental CDR-H3 sequence (GLFDY, hereafter GLF), we constructed several single point mutants (GLH, GLM, GLQ, and ALF) with activity on par with or measurably lower than GLF. We also generated a double mutant (GFA) known to have severely diminished binding activity. When cells expressing these constructs were spot plated under selective conditions, the relative resistance conferred by the five mutants was observed in the following order (from highest to lowest): GLF > GLH ≈ GLM > GLQ ≈ ALF >> GFA (**Fig. 3**). These results were in harmony with the binding activities reported previously for these variants and confirmed that our genetic assay was capable of distinguishing clones on the basis of their relative affinity for antigen.

**Figure 3.**
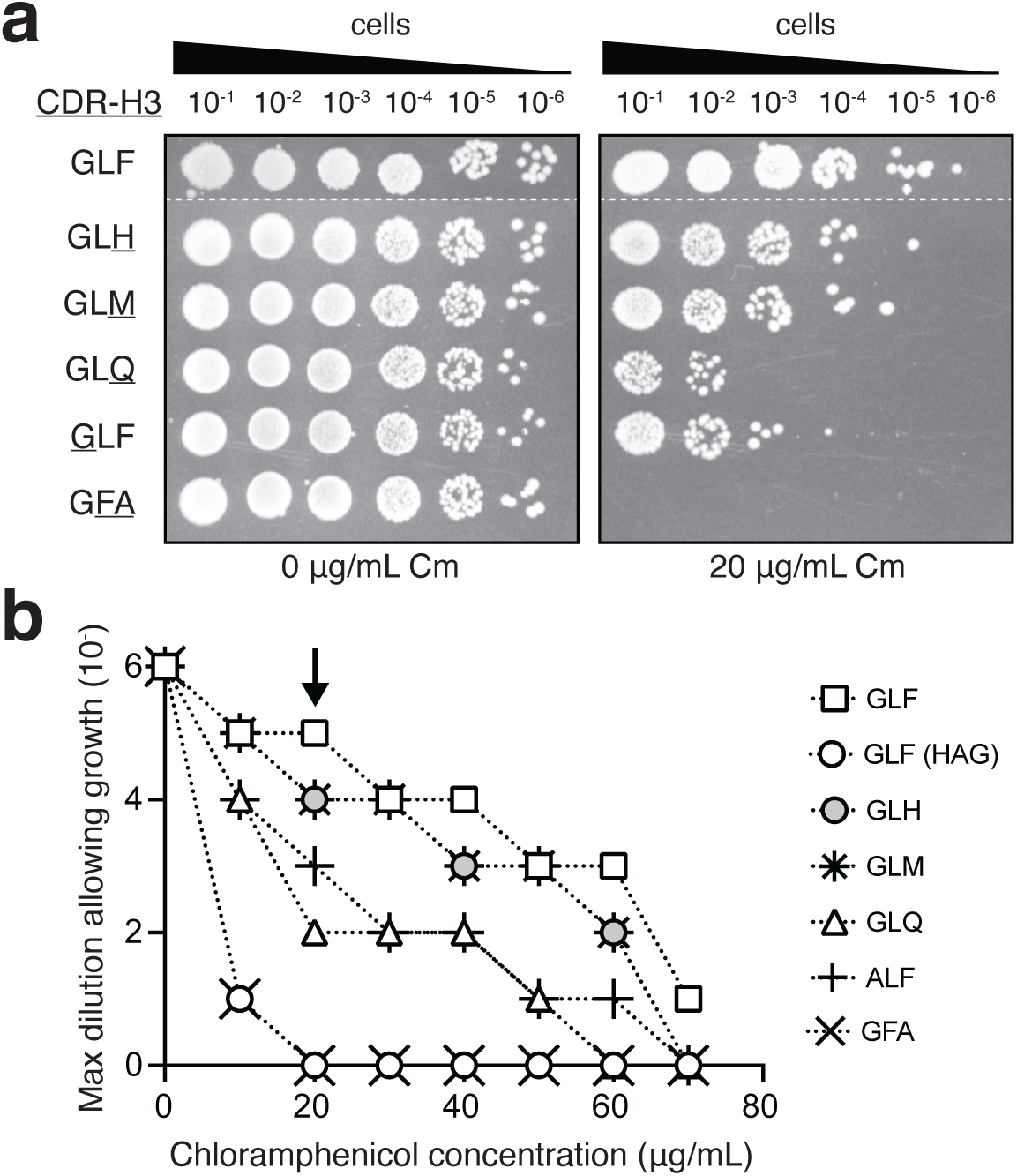
Phenotypic selection of cyclonal variants with differential antigen-binding activity. (a) Representative selective spot plating of SHuffle T7 Express cells carrying a plasmid encoding spTorA-CAT-Gcn4-PP and a second plasmid encoding anti-Gcn4 cyclonal parent (GLF) or variant with CDR-H3 mutation as indicated at left. A total of 5 μl of 10-fold serial diluted cells was plated on LB-agar supplemented with 0 or 20 μg/ml chloramphenicol (Cm) as well as 0.4 % arabinose and 1 mM IPTG to induce protein expression. Spot plating results are representative of at least three biological replicates. Dashed white lines indicate spot plating data merged from discontinuous region of plate. (b) Survival curves for serially diluted SHuffle T7 Express cells co-expressing an anti-Gcn4 cyclonal variant along with the spTorA-CAT-Gcn4-PP reporter. Cells expressing the parental GLF cyclonal along with the non-cognate spTorA-CAT-HAG chimeric antigen (open circle) served as a negative control. Overnight cultures were serially diluted in liquid LB and plated on LB-agar supplemented with Cm. Maximal cell dilution that allowed growth is plotted versus Cm concentration. Arrow in (b) indicates data depicted in image panel (a) and corresponds to 20 μg/ml Cm.

### Selection of novel CDR-H3 cyclonal variants from combinatorial libraries

Encouraged by these results, we next tested whether our selection strategy could be exploited to directly isolate additional Gcn4-PP binders by screening a combinatorial library of cyclonal variants. We chose to randomize the heavy-chain CDR3 based on the fact that this V_H_ region is crucial for determining the specificity for most antibodies ^34^. Indeed, CDRH3 contributes important contacts to the antigen as seen in the crystal structure of the anti-Gcn4 scFv in complex with a Gcn4-derived peptide ^32^. Using the weak-binding GFA cyclonal variant as scaffold, we constructed a library in which the first three residues of CDR-H3 (GFA) were randomized using degenerate codon mutagenesis while the last two (DY) were held constant. Noting that CDR-H3 sequences frequently vary in length, we also constructed a second library based on GFA but with 4 fully randomized positions within a 6-residue heavy-chain CDR3 sequence. The last two residues (DY) were again kept constant.

SHuffle T7 Express cells carrying the plasmid encoding spTorA-CAT-Gcn4-PP were transformed with the cyclonal libraries, after which a total of ∼3×10^7^ clones from each library were selected on agar plates supplemented with 20 μg/ml chloramphenicol. As a negative control, SHuffle T7 Express cells carrying the plasmid encoding spTorA-CAT-Gcn4-PP along with a plasmid encoding the GFA cyclonal variant were plated similarly. After three days, >1,500 colonies appeared on the library plates, while no colonies were observed on the control plates. A total of 20 positive hits were randomly chosen from plates corresponding to each library, and plasmids from all 40 were isolated and retransformed into the same reporter strain to confirm antigen-dependent resistance phenotypes. This test showed that 85% of the originally isolated clones conferred a growth advantage to freshly transformed SHuffle T7 Express cells carrying the chimeric antigen plasmid. Sequencing of the heavy-chain CDR3 region of these positive clones revealed a majority of sequence motifs that were identified in earlier studies (*e*.*g*., GLF, GLH, GLM) ^27,32^; however, three novel motifs were also isolated: GIN, GTK, SLF from the 3-residue library and GLLD from the 4-residue library. Whereas GIN and GLLD were similar to the GIM and GLL motifs that we identified previously ^27^, GTK and SLF were notably different. That is, all 14 unique CDR-H3 sequences reported to date contain only G or A in the first position (with a strong preference for G) and L/V/I in the second position (with a strong preference for L). While there is much weaker conservation in the third position, K has not been observed.

To investigate the binding specificity of these novel CDR-H3s, enzyme-linked immunosorbent assay (ELISA) experiments were carried out with purified versions of Gcn4-PP, HAG, and c-Myc antigens as well as the leucine zipper of the c-Jun proto-oncogene product, which is structurally related to the Gcn4 leucine zipper but not recognized by any anti-Gcn4 scFv intrabodies ^32^. Importantly, all tested cyclonals were highly specific for the cognate Gcn4-PP antigen and did not interact any of the other antigens (**Fig. 4**). In light of these relative binding activities and the close relationship of these motifs to the parental GLF sequence, we conclude that a functional selection for antigen binding in the cytoplasm of *E. coli* has indeed occurred.

**Figure 4.**
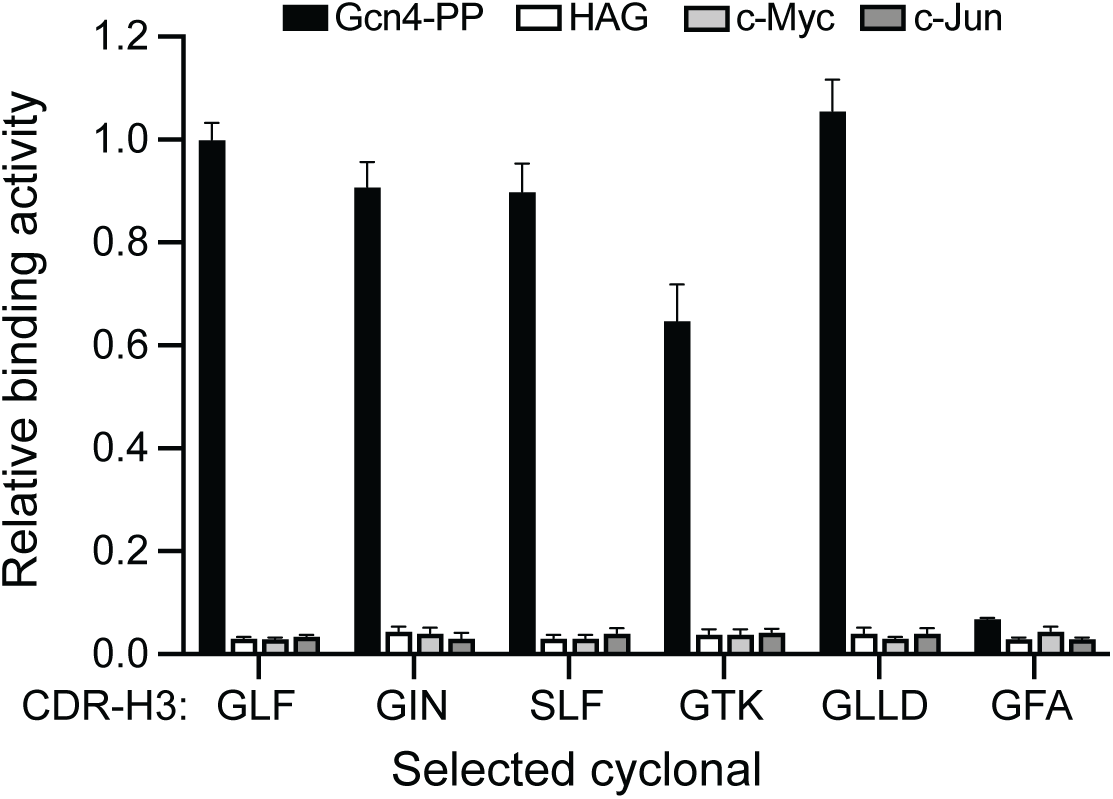
Binding specificity analysis of selected cyclonals by ELISA. (a) Novel cyclonals derived from genetic selection with Gcn4-PP as target antigen were evaluated for interaction with purified GST-Gcn4-PP, GST-HAG, GST-c-Myc, and GST-c-Jun. GLF and GFA cyclonals served as positive and negative controls, respectively. Absorbance was measured at 492 nm and values for each CDR-H3 variant were normalized to the value obtained for GLF. Date are the average of three replicates and error bars represent the standard error of the mean.

## Discussion

In this study, we describe the feasibility of a genetic selection for rapid and reliable isolation of full-length IgG antibodies from combinatorial libraries expressed in the cytoplasm of *E. coli*. This is significant in light of the tremendous impact that recombinant antibodies have made on biomedical research, and increasingly on molecular medicine. Indeed, straightforward technologies that aid in the discovery of mAbs for clinical and therapeutic development remain in high demand. To this end, we designed and validated a high-throughput assay that effectively linked the binding activity of recombinantly expressed IgG antibodies called cyclonals with antibiotic resistance conferred by capture of engineered chimeric antigen biosensors. Using a set of cyclonal variants with the same specificity for one epitope, the leucine zipper domain of yeast Gcn4, we showed that this assay could discriminate antigen-specific cyclonals based on their relative affinities. That is, cells carrying plasmids encoding specific antigen-antibody pairs exhibited an observable fitness advantage over cells carrying plasmids encoding non-specific pairs. The utility of this approach was subsequently revealed by library-based enrichment of several novel anti-GCN4 cyclonal antibodies from a library of randomized CDR-H3 sequences.

Importantly, the results presented here provide the first demonstration of bacterial genetic selection applied to the discovery of full-length IgG antibodies. Genetic selections are attractive as they link a desired property, in this case antigen-binding activity, to the fitness of the host organism. To date, a handful of genetic selections have been reported for isolating functional antibodies in bacteria and yeast; however, these have only been demonstrated for scFvs and other small formats ^25,27,35-41^ but not full-length IgGs. Indeed, the vast majority of recombinant antibody screening platforms in microorganisms make use of scFv or Fab antibodies ^8^. While these formats are relatively easy to produce in bacteria and yeast, they are monovalent proteins that typically lack avidity effects which can be important for reducing antigen off-rates and for enhancing the recovery of low-affinity binders ^42^. Moreover, these monovalent formats are generally unsuitable for therapeutic development and must be converted to full-length IgGs prior to use in the clinic. Unfortunately, the conversion process requires additional cloning steps and can result in loss of binding activity ^12^. By leveraging full-length IgG expression in the bacterial cytoplasm ^19^, our approach obviates the need for post-selection molecular reformatting.

Another advantage of our approach is that genetic selection is intrinsically high throughput, enabling cyclonal variants with desirable binding activity to be readily isolated from large libraries by simple transformation and plating of bacteria without needing to purify or immobilize the target antigen. While not directly demonstrated here, our selection strategy should permit selection of very large libraries (>10^11^). Screens, on the other hand, require every member of a library to be analyzed, making the process of identifying clones with beneficial mutations much more labor intensive. For example, many of the previous display-based methods involve fluorescence activated cell sorting (FACS), a very powerful high-throughput screening methodology; however, interrogating a library of >10^8^ cells using FACS is time-consuming and technically challenging ^12,17^. In fact, for combinatorial libraries of this size, an initial phage display screening process was required to reduce the initial library to a size that was manageable by FACS ^12^.

A final advantage is that unlike nearly all other full-length IgG screening methods that require tethering of the antibody to a cellular membrane, using either fusion to a membrane anchoring polypeptide ^15,16^ or introduction of a secretion-and-capture step prior to antigen binding ^10-14^, our ‘membrane-less’ approach does not depend on physical display of the antibody. In fact, our method of cytoplasmic IgG expression circumvents membrane translocation of these large macromolecules altogether, which is important because traversing tightly sealed biological membranes is a rate limiting and energy intensive step that can serve as a potential source of selection bias in these previous IgG screening methods. While other membrane-less IgG screening strategies exist, in particular methods for encapsulating single IgG antibody secreting cells in water-in-oil droplets ^43,44^ or gel microdroplets ^45-48^, construction of such drop-based secretor cell libraries is non-trivial, often involving microfluidics, and screening must typically be performed in conjunction with FACS, which introduces additional challenges as discussed above. It should also be pointed out that because our selection requires no modification of the IgG, the selected plasmid can be used directly for functional IgG expression without any subcloning, thereby streamlining the process from selection to expression of IgG antibody products.

In conclusion, we have demonstrated a promising new methodology for stringent selection of full-length IgG antibodies from combinatorial libraries with the potential to yield high-affinity binders with selective target binding characteristics. In the future, we anticipate that this system will find use in the isolation of entirely new antibodies by functionally interrogating more complex libraries comprised of naïve antibody repertoires as well as in the engineering of ultra-high affinity IgG antibodies by affinity maturing parental antibody sequences using directed evolution workflows. With these and other imagined uses, our recombinant antibody selection technology represents a powerful new addition to the antibody engineering toolkit that should facilitate discovery of antibody-based research reagents, diagnostics, and biopharmaceuticals in the years to come.

## Materials and Methods

### Bacterial strains

*E. coli* strain DH5α was used for plasmid construction while SHuffle T7 Express (New England Biolabs) ^20^ was used for cyclonal expression and library selections. Protein antigens for immunoassays including GST-Gcn4-PP, GST-HAG, GST-c-Myc, and GST-c-Jun were expressed using *E. coli* T7 Express (New England Biolabs).

### Plasmid construction

The pBAD24 plasmid ^49^ was used for construction of all spTorA-CAT-Ag chimeric antigen reporter fusions. First, a PCR product corresponding to spTorA-JunLZ-FLAG ^27^, encoding the signal peptide of *E. coli* TorA (spTorA) fused to the N-terminus of the c-Jun leucine zipper (JunLZ) was cloned between the NheI and HindIII restriction sites of pBAD24, yielding plasmid pBAD24-spTorA-JunLZ-FLAG. Next, the gene encoding CAT was PCR-amplified from pACYC-Duet™-1 (Novagen) to include a 3’ flexible linker (GTSAAAG) flanked by SalI and SpeI restriction sites. At the same time, the gene encoding Gcn4(7P14P) ^27,32^, encoding a double proline mutant of the leucine zipper domain of Gcn4 that reduces its propensity for homodimerization, was PCR-amplified from pBAD33-Gcn4(7P14P)-Bla ^27^ to include the same flexible linker sequence at the 5’ end. The two resulting PCR products were fused by overlap extension PCR and the overlap product was cloned between the XbaI and HindIII sites of pBAD24-spTorA-JunLZ-FLAG, yielding plasmid pBAD24-spTorA-CAT-Gcn4-PP. Genes encoding the HAG (DVPDYA) and c-Myc (EQKLISEEDL) epitopes were constructed by annealing complementary oligonucleotides, and were subsequently cloned in place of Gcn4(7P14P) between SpeI and HindIII sites in pBAD24-spTorA-CAT-Gcn4-PP, yielding plasmids pBAD24-spTorA-CAT-HAG and pBAD24-spTorA-CAT-c-Myc.

The creation of all bacterial IgG expression constructs involved plasmid pCOLADuet™-1 (Novagen), which is designed for the coexpression of two target genes from independent upstream T7 promoter/lac operator regions. First, the light chain genes (V_L_-mC_L_κ) for anti-HAG and anti-Gcn4 were PCR-amplified from pMAZ360-cIgG-aHAG and pMAZ360-cIgG-aGcn4 ^19^, respectively, and cloned between NcoI and NotI sites of pCOLA-Duet™-1, yielding plasmids pCD1-cLC-aHAG and pCD1-cLC-aGcn4, respectively. Next, the heavy chain Fab genes (V_H_-mC_H_1) were PCR-amplified from the same pMAZ360 templates and cloned between NdeI and AscI sites in pCD1-cLC-aHAG and pCD1-cLC-aGcn4, yielding plasmids pCD1-cFab-aHAG and pCD1-cFab-aGcn4. Finally, the heavy chain Fc genes (hFc) were PCR-amplified from the pMAZ360 template plasmids and cloned between AscI and XhoI sites in pCD1-cFab-aHAG and pCD1-cFab-aGcn4, yielding plasmids pCD1-cIgG-aHAG and pCD1-cIgG-aGcn4.

To construct the anti-c-Myc cyclonal, the gene encoding the V_L_ domain of scFv-3DX ^31^ was PCR-amplified using primers that introduced a sequence overlapping with the mouse constant light chain kappa domain (mC_L_κ). In parallel, the gene encoding mCLκ was PCR-amplified with primers that introduced a 5’ sequence overlapping with the V_L_ of scFv-3DX. The resulting PCR products were assembled by overlap extension PCR, generating the anti-c-Myc light chain (V_L_-mC_L_κ). Similarly, the gene encoding the V_H_ domain of scFv-3DX was PCR-amplified using primers that introduced a sequence overlapping with the mFab/hFc heavy chain constant domains. At the same time, the mFab/hFc constant heavy chain domains were amplified with primers that introduced a 5’ sequence overlapping with V_H_ of scFv-3DX. Again, the resulting products were assembled by overlap extension PCR, generating the anti-c-Myc heavy chain (V_H_-mC_H_1-hFc). The light chain and heavy chain products were then cloned between NcoI/NotI and NdeI/XhoI sites, respectively, of pCOLADuet™-1, yielding the plasmid pCD1-cIgG-c-Myc. The heavy-chain CDR3 cyclonal variants GLH, GLM, GLQ, ALF, and GFA were constructed by site-directed mutagenesis of the parental GLF cyclonal heavy chain sequence. Plasmids pET28a-GST-Gcn4-PP and pET28a-HAG were described previously ^27^. An identical strategy was used to construct pET28a-GST-c-Myc and pET28a-GST-c-Jun. All plasmids constructed in this study were confirmed by sequencing at the Cornell Biotechnology Resource Center.

### Selective growth assays

Chemically competent SHuffle T7 Express cells were transformed with one of the pBAD24-spTorA-CAT-Ag plasmids along with a pCD1-cyclonal plasmid, and spread on Luria-Bertani (LB)-agar plates supplemented with 25 μg/ml spectinomycin (Spec), 25 μg/ml kanamycin (Kan), and 50 μg/ml ampicillin (Amp), and cultured overnight at 37°C. The next day, 3 mL of LB supplemented with appropriate antibiotics was inoculated with three freshly transformed colonies and incubated at 30°C for 12-18 h. Cells carrying the pBAD24-spTorA-CAT-Ag and pCD1-cyclonal plasmids were normalized to an absorbance at 600 nm (Abs_600_) ≈ 2.5 (2.5×10^9^ cells/mL). Cells were then serially diluted ten-fold in liquid LB, and 5 μl of each dilution was spotted on selective induction plates supplemented with 25 μg/ml Spec, 25 μg/ml Kan, 50 μg/ml Amp, 1 M IPTG, 0.2% (w/v) arabinose, and varying concentrations of Cm. The plates were then incubated at 30°C for 24-48 h.

### Library construction

Random mutagenesis of the first three residues of CDR-H3 was performed using NDT and NNK degenerate codons. The resulting library encoded anti-Gcn4 cyclonals with heavy-chain CDR3 motifs of the form XXXDY, where X was encoded by either the NDT codon (encoding 12 amino acids: N, S, I, H, R, L, Y, C, F, D, G, and V; and no stop codons) or NNK (encoding all amino acids and one stop codon. Random mutagenesis of the CDR-H3 was achieved by amplifying the entire pCD1-cIgG-aGcn4(GFA) plasmid by inverse PCR with degenerate NDT and NNK primers encoding the three randomized codons within CDR-H3. The resulting linear PCR product was circularized by blunt-end ligation to produce the plasmid library. The circularized products were used to transform electrocompetent DH5α cells. The transformed cells were cultured overnight in 100 mL LB supplemented with 50 μg/ml Kan. Plasmid DNA was purified by maxiprep from the overnight culture for selection experiments. Random mutagenesis of the first four residues of CDR-H3 was performed identically using a degenerate NDT primer to generate six-residue heavy-chain CDR3 motifs of the form XXXXDY.

### Library selection

To perform library selections, electrocompetent SHuffle T7 Express cells carrying pBAD24-spTorA-CAT-Gcn4-PP were transformed with the purified anti-Gcn4 cyclonal libraries. Transformants were incubated in SOC media at 37°C for 1 h without antibiotics and then cultured overnight in LB supplemented with appropriate antibiotics and 0.2% glucose. The next day, overnight cells were normalized to Abs_600_ ≈ 2.5 and serially diluted to 10^−3^, 10^−4^, and 10^−5^. A total volume of 225 µl of each dilution was plated on LB-agar supplemented with 15-30 μg/ml Cm, 0.2% (w/v) arabinose, and 1 mM IPTG and cultured at 30°C for 72 h. At the same time, cells transformed with plasmid pBAD24-spTorA-CAT-Gcn4-PP and pCD1-cIgG-Gcn4(GFA) and were treated in an identical manner as library cells and served as a negative control. Clones that appeared on selective plates were picked at random and resistance to Cm was verified by isolating plasmid DNA, retransforming SHuffle T7 Express cells, and performing selective spot plating with the freshly transformed cells. Plasmid DNA of verified positive hits was sequenced at the Cornell Biotechnology Resource Center.

### Preparation of soluble cell extracts and ELISA

A single colony of SHuffle T7 Express carrying one of the pCD1-cyclonal plasmids was used to inoculate 2 ml LB supplemented with appropriate antibiotics, and grown overnight at 30 °C. The next day, 5 ml of fresh LB supplemented with appropriate antibiotics was inoculated 1/100 with the overnight culture and cells were grown at 30 °C until reaching Abs_600_ ≈ 0.7. At this point, cyclonal expression was induced by addition of 0.1 mM IPTG, after which cells were incubated an additional 16 h at RT or 30°C. Cells were harvested by centrifugation before preparation of lysates. Cells expressing recombinant proteins were harvested by centrifugation (4,000 x g, 4°C) and resuspended in PBS and 5 mM EDTA. Cells were lysed in an ice-water bath by sonication (Branson sonifier 450; duty cycle 30%, output control 3) using four repetitions of 30 s each. The insoluble fraction was removed by centrifugation (21,000 x g, 4°C) and the supernatant was collected as the soluble fraction.

The GST-Gcn4-PP, GST-HAG, GST-c-Myc, and GST-c-Jun fusion proteins were expressed in *E. coli* T7 express cells and purified using Ni-NTA affinity resin according to standard protocols. Next, Costar 96-well ELISA plates (Corning) were coated overnight at 4°C with 50 µl of 10 µg/ml of each of the different GST fusions, in 0.05 M sodium carbonate buffer (pH 9.6). After blocking in PBST with 3% (w/v) milk (PBSTM) for 1–3 h at room temperature, the plates were washed four times with PBS buffer and incubated with serially diluted soluble fractions of crude cell lysates for 1 h at room temperature. Cyclonal IgG-containing samples were quantified by the Bradford assay and an equivalent amount of total protein (typically 8–64 mg) was applied to the plate. After washing four times with the same buffer, 50 µl of 1:5,000-diluted rabbit anti-human IgG (Fc) antibody–HRP conjugate (Pierce) antibodies in PBSTM was added to each well for 1 h. The 96-well plates were then washed six times with PBST. After the final wash, 200 µl SigmaFAST(tm) OPD solution (Sigma-Aldrich) was added and incubated in each well in the dark for 30 min. The HRP reaction was then terminated by the addition of 50 µl 3 M H_2_SO_4_ to the wells. Following reaction quenching, the absorbance of each well was measured at 492 nm.

## Acknowledgements

This work was supported by the National Institutes of Health grant #AI092969-01A1 (to M.P.D. and M.B.), the Defense Threat Reduction Agency (GRANT11631647 to M.PD.), the National Science Foundation (CBET-1605242 to M.P.D.), the National Cancer Institute (U54 CA210184 seed project to M.P.D.), a Ford Foundation Predoctoral Fellowship (to M.-P.R.), a National Science Foundation Graduate Research Fellowship (to M.-P.R.), an NIH Chemical-Biology Interface (CBI) training fellowship (supporting grant T32GM008500 to E.C.C.), and a Royal Thai Government Graduate Fellowship (to T.J.).

## Author contributions

M.-P.R. designed and performed research, analyzed data, and wrote the paper. E.C.C., M.L., T.J., X.Z. and M.Z. performed research. M.B. and M.P.D. conceptualized the project, designed research, analyzed data, and wrote the paper.

## References

1. Kohler, G. & Milstein, C. Continuous cultures of fused cells secreting antibody of predefined specificity. Nature 256, 495–7 (1975).

2. Li, J. et al. Human antibodies for immunotherapy development generated via a human B cell hybridoma technology. Proc Natl Acad Sci U S A 103, 3557–62 (2006).

3. Fishwild, D.M. et al. High-avidity human IgG kappa monoclonal antibodies from a novel strain of minilocus transgenic mice. Nat Biotechnol 14, 845–51 (1996).

4. Winter, G., Griffiths, A.D., Hawkins, R.E. & Hoogenboom, H.R. Making antibodies by phage display technology. Annu Rev Immunol 12, 433–55 (1994).

5. Lipovsek, D. & Pluckthun, A. In-vitro protein evolution by ribosome display and mRNA display. J Immunol Methods 290, 51–67 (2004).

6. Daugherty, P.S., Chen, G., Olsen, M.J., Iverson, B.L. & Georgiou, G. Atntibody affinity maturation using bacterial surface display. Protein Eng 11, 825–32 (1998).

7. Harvey, B.R. et al. Anchored periplasmic expression, a versatile technology for the isolation of high-affinity antibodies from Escherichia coli-expressed libraries. Proc Natl Acad Sci U S A 101, 9193–8 (2004).

8. Hoogenboom, H.R. Selecting and screening recombinant antibody libraries. Nat Biotechnol 23, 1105–16 (2005).

9. Boder, E.T. & Wittrup, K.D. Yeast surface display for screening combinatorial polypeptide libraries. Nat Biotechnol 15, 553–7 (1997).

10. Mazor, Y., Van Blarcom, T., Mabry, R., Iverson, B.L. & Georgiou, G. Isolation of engineered, full-length antibodies from libraries expressed in Escherichia coli. Nat Biotechnol 25, 563–5 (2007).

11. Mazor, Y., Van Blarcom, T., Iverson, B.L. & Georgiou, G. E-clonal antibodies: selection of full-length IgG antibodies using bacterial periplasmic display. Nat Protoc 3, 1766–77 (2008).

12. Mazor, Y., Van Blarcom, T., Carroll, S. & Georgiou, G. Selection of full-length IgGs by tandem display on filamentous phage particles and Escherichia coli fluorescence-activated cell sorting screening. The FEBS journal 277, 2291–2303 (2010).

13. Rakestraw, J.A., Aird, D., Aha, P.M., Baynes, B.M. & Lipovsek, D. Secretion-and-capture cell-surface display for selection of target-binding proteins. Protein Eng Des Sel 24, 525–30 (2011).

14. Rhiel, L. et al. REAL-Select: full-length antibody display and library screening by surface capture on yeast cells. PLoS One 9, e114887 (2014).

15. Shaheen, H.H. et al. A dual-mode surface display system for the maturation and production of monoclonal antibodies in glyco-engineered Pichia pastoris. PLoS One 8, e70190 (2013).

16. Zhou, C., Jacobsen, F.W., Cai, L., Chen, Q. & Shen, W.D. Development of a novel mammalian cell surface antibody display platform. MAbs 2, 508–18 (2010).

17. Georgiou, G. Analysis of large libraries of protein mutants using flow cytometry. Adv Protein Chem 55, 293–315 (2000).

18. Lee, P.A., Tullman-Ercek, D. & Georgiou, G. The bacterial twin-arginine translocation pathway. Annu Rev Microbiol 60, 373–95 (2006).

19. Robinson, M.P. et al. Efficient expression of full-length antibodies in the cytoplasm of engineered bacteria. Nat Commun 6, 8072 (2015).

20. Lobstein, J. et al. SHuffle, a novel Escherichia coli protein expression strain capable of correctly folding disulfide bonded proteins in its cytoplasm. Microb Cell Fact 11, 56 (2012).

21. DeLisa, M.P., Tullman, D. & Georgiou, G. Folding quality control in the export of proteins by the bacterial twin-arginine translocation pathway. Proc Natl Acad Sci U S A 100, 6115–20 (2003).

22. Fisher, A.C., Kim, W. & DeLisa, M.P. Genetic selection for protein solubility enabled by the folding quality control feature of the twin-arginine translocation pathway. Protein Sci 15, 449–58 (2006).

23. Stanley, N.R. et al. Behaviour of topological marker proteins targeted to the Tat protein transport pathway. Mol Microbiol 43, 1005–21 (2002).

24. Hicks, M.G., Lee, P.A., Georgiou, G., Berks, B.C. & Palmer, T. Positive selection for loss-of-function tat mutations identifies critical residues required for TatA activity. J Bacteriol 187, 2920–5 (2005).

25. Lee, H.C., Portnoff, A.D., Rocco, M.A. & DeLisa, M.P. An engineered genetic selection for ternary protein complexes inspired by a natural three-component hitchhiker mechanism. Sci Rep 4, 7570 (2014).

26. Rodrigue, A., Chanal, A., Beck, K., Muller, M. & Wu, L.F. Co-translocation of a periplasmic enzyme complex by a hitchhiker mechanism through the bacterial tat pathway. J Biol Chem 274, 13223–8 (1999).

27. Waraho, D. & DeLisa, M.P. Versatile selection technology for intracellular protein-protein interactions mediated by a unique bacterial hitchhiker transport mechanism. Proc Natl Acad Sci U S A 106, 3692–7 (2009).

28. Otrelo-Cardoso, A.R. et al. Structural data on the periplasmic aldehyde oxidoreductase PaoABC from Escherichia coli: SAXS and preliminary X-ray crystallography analysis. Int J Mol Sci 15, 2223–36 (2014).

29. Rayner, L.E. et al. The solution structures of two human IgG1 antibodies show conformational stability and accommodate their C1q and FcgammaR ligands. J Biol Chem 290, 8420–38 (2015).

30. Jermutus, L., Honegger, A., Schwesinger, F., Hanes, J. & Pluckthun, A. Tailoring in vitro evolution for protein affinity or stability. Proc Natl Acad Sci U S A 98, 75–80 (2001).

31. Fujiwara, K. et al. A single-chain antibody/epitope system for functional analysis of protein-protein interactions. Biochemistry 41, 12729–38 (2002).

32. der Maur, A.A. et al. Direct in vivo screening of intrabody libraries constructed on a highly stable single-chain framework. J Biol Chem 277, 45075–85 (2002).

33. Cristobal, S., de Gier, J.W., Nielsen, H. & von Heijne, G. Competition between Sec- and TAT-dependent protein translocation in Escherichia coli. EMBO J 18, 2982–90 (1999).

34. Xu, J.L. & Davis, M.M. Diversity in the CDR3 region of V(H) is sufficient for most antibody specificities. Immunity 13, 37–45 (2000).

35. Waraho-Zhmayev, D., Meksiriporn, B., Portnoff, A.D. & DeLisa, M.P. Optimizing recombinant antibodies for intracellular function using hitchhiker-mediated survival selection. Protein Eng Des Sel 27, 351–8 (2014).

36. Meksiriporn, B. et al. A survival selection strategy for engineering synthetic binding proteins that specifically recognize post-translationally phosphorylated proteins. Nat Commun 10, 1830 (2019).

37. Koch, H., Grafe, N., Schiess, R. & Pluckthun, A. Direct selection of antibodies from complex libraries with the protein fragment complementation assay. J Mol Biol 357, 427–41 (2006).

38. Mossner, E., Koch, H. & Pluckthun, A. Fast selection of antibodies without antigen purification: adaptation of the protein fragment complementation assay to select antigen-antibody pairs. J Mol Biol 308, 115–22 (2001).

39. Lofdahl, P.A., Nord, O., Janzon, L. & Nygren, P.A. Selection of TNF-alpha binding affibody molecules using a beta-lactamase protein fragment complementation assay. N Biotechnol 26, 251–9 (2009).

40. Lofdahl, P.A. & Nygren, P.A. Affinity maturation of a TNFalpha-binding affibody molecule by Darwinian survival selection. Biotechnol Appl Biochem 55, 111–20 (2010).

41. Secco, P. et al. Antibody library selection by the {beta}-lactamase protein fragment complementation assay. Protein Eng Des Sel 22, 149–58 (2009).

42. Lee, C.V., Sidhu, S.S. & Fuh, G. Bivalent antibody phage display mimics natural immunoglobulin. J Immunol Methods 284, 119–32 (2004).

43. El Debs, B., Utharala, R., Balyasnikova, I.V., Griffiths, A.D. & Merten, C.A. Functional single-cell hybridoma screening using droplet-based microfluidics. Proc Natl Acad Sci U S A 109, 11570–5 (2012).

44. Koster, S. et al. Drop-based microfluidic devices for encapsulation of single cells. Lab Chip 8, 1110–5 (2008).

45. Akbari, S. & Pirbodaghi, T. A droplet-based heterogeneous immunoassay for screening single cells secreting antigen-specific antibodies. Lab Chip 14, 3275–80 (2014).

46. Fang, Y., Chu, T.H., Ackerman, M.E. & Griswold, K.E. Going native: Direct high throughput screening of secreted full-length IgG antibodies against cell membrane proteins. MAbs 9, 1253–1261 (2017).

47. Mettler Izquierdo, S. et al. High-efficiency antibody discovery achieved with multiplexed microscopy. Microscopy (Oxf) 65, 341–52 (2016).

48. Powell, K.T. & Weaver, J.C. Gel microdroplets and flow cytometry: rapid determination of antibody secretion by individual cells within a cell population. Biotechnology (N Y) 8, 333–7 (1990).

49. Guzman, L.M., Belin, D., Carson, M.J. & Beckwith, J. Tight regulation, modulation, and high-level expression by vectors containing the arabinose PBAD promoter. J Bacteriol 177, 4121–30 (1995).

